# Human HelQ and DNA polymerase δ interact to halt DNA synthesis and stimulate DNA single-strand annealing

**DOI:** 10.1101/2022.10.02.510506

**Authors:** Liu He, Rebecca Lever, Andrew Cubbon, Muhammad Tehseen, Tabitha Jenkins, Alice O. Nottingham, Anya Horton, Hannah Betts, Martin Fisher, Samir M. Hamdan, Panos Soultanas, Edward L. Bolt

## Abstract

DNA strand breaks can be repaired by base-pairing with unbroken homologous DNA, forming a template for new DNA synthesis that patches over the break site. In eukaryotes multiple DNA break repair pathways utilize DNA polymerase δ (Pol δ) to synthesise new DNA from the available 3’OH at the strand break. Here we show that DNA synthesis by human Pol δ is halted by the HelQ DNA repair protein directly targeting isolated Pol δ or Pol δ in complex with PCNA and RPA. The mechanism is independent of DNA binding by HelQ or Pol δ, maps to a region of HelQ that also modulates RPA, and requires multiple Pol δ subunits. Interaction of HelQ with the POLD3 subunit of Pol δ stimulated DNA single-strand annealing activity of HelQ. The data implicates HelQ in preventing genetic instability by restraining DNA synthesis in multiple DNA break repair pathways.

## Introduction

DNA strand breaks are repaired by homologous recombination or micro-homology mediated end-joining reliant on new DNA synthesis that patches over the broken region. DNA polymerases synthesise DNA in these contexts by extending D-loops generated by a recombinase Rad51/RecA/RadA or from base-paired DNA microhomologies [1-8]. In eukaryotes DNA polymerase δ (Pol δ) synthesises DNA during multiple modes of homologous recombination that trigger varying genetic outcomes according to the context of recombination and the extent of DNA synthesis–in DNA break repair by Break-Induced Replication (BIR) or Microhomology-Mediated BIR (MM-BIR) DNA replication triggers genetic rearrangements, tandem duplications and mutagenesis characteristic of cells coping with chronic DNA damage and replication stress [4, 9-12]. Mechanisms have evolved that limit or prevent mutagenic DNA synthesis during HR.

In eukaryotes helicase enzymes conserved from yeasts to humans prevent or dissociate D-loop DNA structures, antagonising the priming of new DNA synthesis from a strand break [13-19]. The DNA helicase HelQ contributes to DNA break repair in metazoans but it is absent from yeasts. Loss of HelQ predisposes to cancers and infertility [20-23] and corresponds to increased frequencies of tandem DNA duplications and long-tracts of DNA repair synthesis during homologous recombination [24, 25], and to reduced repair by SDSA [26]. HelQ promotes DNA strand annealing in contexts distinct from microhomology-mediated DNA repair by its homologue PolQ, a polymerase-helicase that is also found only in metazoans [24, 25]. The opposing HelQ helicase unwinding and annealing activities may be balanced through its interaction with RPA that stimulates annealing [25, 27], and with Rad51 that stimulates HelQ helicase activity [25]. It is not clear how these activities of HelQ may limit DNA replication to shorter tracts that prevent mutagenesis in the form of DNA duplications. Here we show that human HelQ halts DNA synthesis by direct targeting of Pol δ, and that this stimulates HelQ to anneal homologous single-stranded DNA. This identifies a plausible mechanism in which HelQ restrains DNA synthesis during homologous recombination.

## Results

### Targeting of DNA polymerase δ by HelQ halts DNA synthesis

We investigated if HelQ modulates DNA synthesis reactions *in vitro* as a possible explanation for characteristic genetic instability caused by its loss [24]. Purified HelQ protein (Figure S1A) was introduced into primer extension reactions, including catalysed by Pol δ. Purified human Pol δ comprising four subunits (Figure S1A, POLD1/p125, POLD2/p50, POLD3/p68 and POLD4/p12 [28]) in complex with PCNA and RPA catalyses processive DNA synthesis. In our assays this was apparent from Pol δ (100 nM) being unable to synthesise new DNA from a 25nt Cy5 labelled primer-M13 template unless it was pre-incubated with purified PCNA and RPA, summarised in Figure 1A (lanes 1-3 and 5-7). Synthetic products from the Pol δ-RPA-PCNA holoenzyme, visible by ethidium bromide staining and imaging of Cy5, were severely reduced by adding HelQ (100 nM) to reactions after Pol δ had been pre-incubated with RPA-PCNA (Figure 1A lanes 4 and 8). HelQ removes RPA from ssDNA [25, 27], providing a potential explanation for its inhibitory effect on primer extension if RPA were no longer available to Pol δ. Therefore we switched to Pol δ primer extension reactions that synthesise DNA from a primer annealed to a short oligonucleotide template without need for RPA and PCNA (Figure1B and S1B). Again we observed inhibition of Pol δ (40 nM) to zero detectable product formation on titrating HelQ (40-160 nM) into the reactions immediately after addition of Pol δ to DNA (Figure 1C).

**Figure 1:**
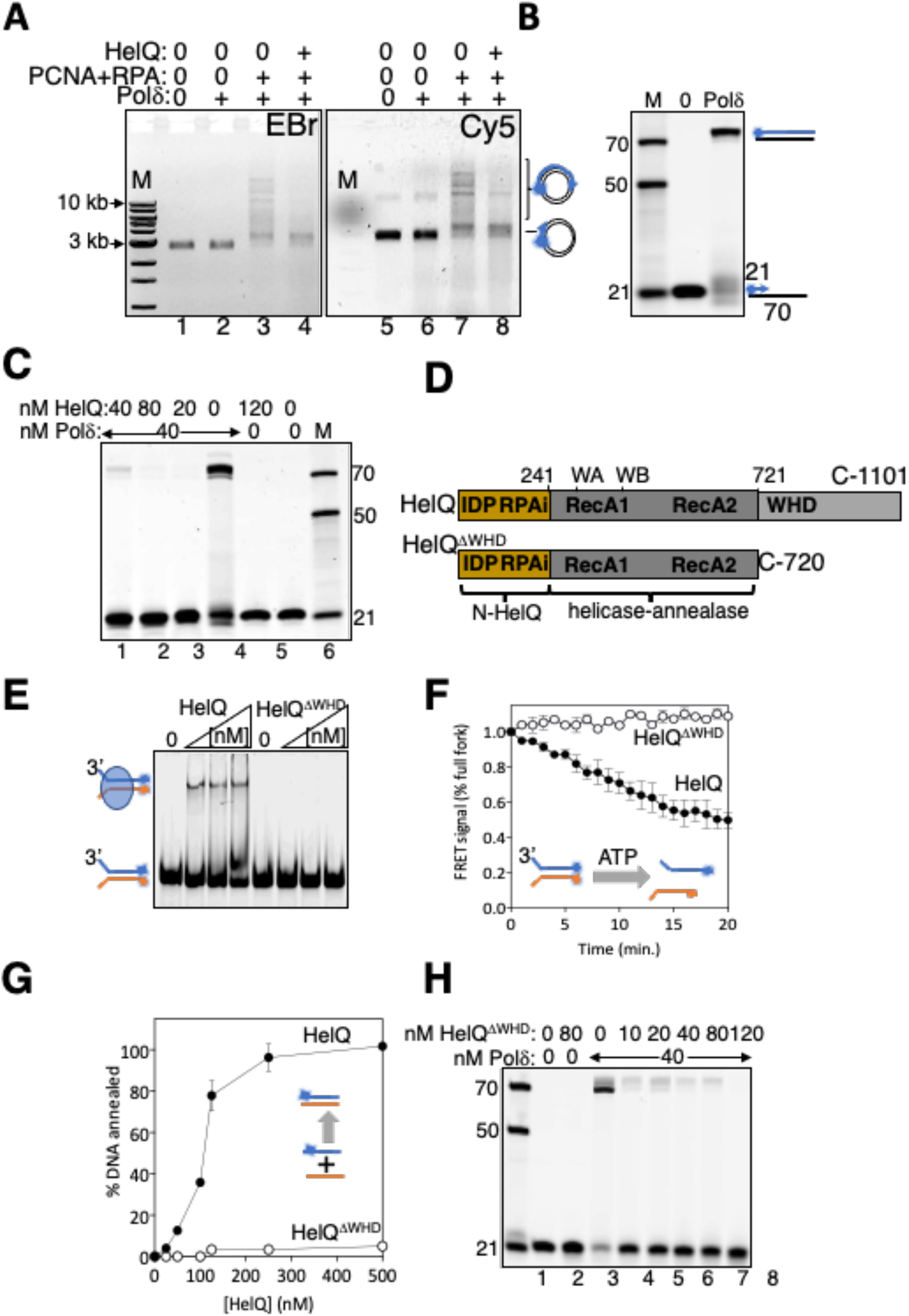
**(A)**. Primer extension by Pol δ-RPA-PCNA-RFC holoenzyme is inhibited by addition of HelQ, summarised in an ethidium bromide-stained agarose gel (EBr, lanes 1-4) and by imaging extended Cy5-labelled (blue circle) oligonucleotides from the same gel (Cy5, lanes 5-8). Proteins present in reactions are indicated (+); Pol δ (40 nM), PCNA (40 nM), RPA (320 nM), RFC (12 nM) and HelQ (100 nM) were added to M13 DNA (1.32 nM). DNA marker (M) sizes are indicated. **(B)**. Pol δ alone (40 nM) catalyses extension of a Cy5 (blue circle) end labelled 21nt primer (15 nM) annealed to a 70nt template, visualised by Typhoon imaging of the denaturing (8 M urea) gel. Cy5 end-labelled DNA markers are indicated by length to the left of the panel, (21, 50 and 70nt), and in all subsequent gel panels. To the right of the panel depicts the fully extended product of Pol δ that is visible in the urea gel as the extended cy5-lablled ssDNA. **(C)**. DNA synthesis by Pol δ (40 nM) was inhibited by titration of HelQ, lanes 1-3, compared with uninhibited DNA synthesis (lane 4) and including control reactions (lanes 5 and 6), with proteins included as indicated. **(D)**. For context, HelQ^ΔWHD^ is shown alongside full-length HelQ protein to indicate catalytic domains (RecA1 and A2), the predicted winged helix domain (WHD) [27, 30], and the non-catalytic N-HelQ fragment (see also Figure 2) that comprises an intrinsically disordered protein (IDP) region and a region known to modulate RPA function (RPAi [27]). **(E)**. EMSA showing HelQ^ΔWHD^ (0, 100, 200, 400nM) is unable to detectably bind to a model forked DNA substrate (15 nM) that is favoured for binding and unwinding by HelQ [27] (0, 100, 200, 400nM). **(F)**. HelQ^ΔWHD^ compared with HelQ (both 150 nM) is unable to unwind a model forked DNA substrate (15 nM) measured by decreased FRET efficiency over time from separation of Cy5 and Cy3 labelled DNA strands as shown, **(G)** HelQ^DWHD^ is severely deficient in DNA strand annealing compared with HelQ measured from gel-based assays pairing of complementary DNA strands. **(E)** Urea gel summarising that DNA synthesis by Pol δ (40nM, lane 3) is inhibited by titration of the HelQ^ΔWHD^ at concentrations indicated.

HelQ binding to ssDNA activates its DNA translocation/helicase activity [27, 29]. This provided a further possible explanation for its inhibitory effect on Pol δ, if it formed a physical barrier to primer extension or out-competed Pol δ for access to DNA. To test this we isolated a novel HelQ protein (HelQ^ΔWHD^, Figure 1D and S1A) that lacks a predicted DNA-binding winged helix domain [30]. HelQ^ΔWHD^ was unable to bind to DNA in EMSAs (summarised in Figure 1E) and consistent with this was inactive at DNA unwinding, and only weakly annealed DNA, measured in real time by FRET efficiencies and confirmed in gel products observed from end point assays (Figure 1F and 1G, and S1C and S1D). However, HelQ^ΔWHD^ halted primer extension by Pol δ (Figure 1H). We concluded that HelQ targets Pol δ directly to halt DNA synthesis by a mechanism that is does not require HelQ-RPA or HelQ-DNA interactions.

### A catalytically inactive fragment of HelQ selectively inhibits DNA synthesis by Pol δ

To learn more about the inhibitory mechanism we turned to a 241-amino acid HelQ region at the extreme N-terminus, termed N-HelQ (see Figure 1D) that is unable to bind to DNA and is catalytically inactive [27]. Purified N-HelQ also inhibited Pol δ (40 nM), reducing primer extension to 20% of full activity when at 10 nM and to zero at 120 nM (Figure 2A and 2B), further confirming that neither HelQ-DNA binding or catalytic activity is required for inhibiting primer extension by Pol δ. Consistent with this, the inhibitory effect of N-HelQ was enhanced at least 5-fold by pre-mixing it with Pol δ for 5 minutes before adding DNA to reactions (1-16 nM N-HelQ: 40 nM Pol δ) (Figure 2C, 2D and S2A). N-HelQ was also selective at inhibiting DNA synthesis by Pol δ – human polymerase η (Pol η) and polymerase κ (Pol κ), and bacterial polymerases I and III, were all unaffected by addition of N-HelQ to primer extension reactions (Figure 2E and S2B).

**Figure 2:**
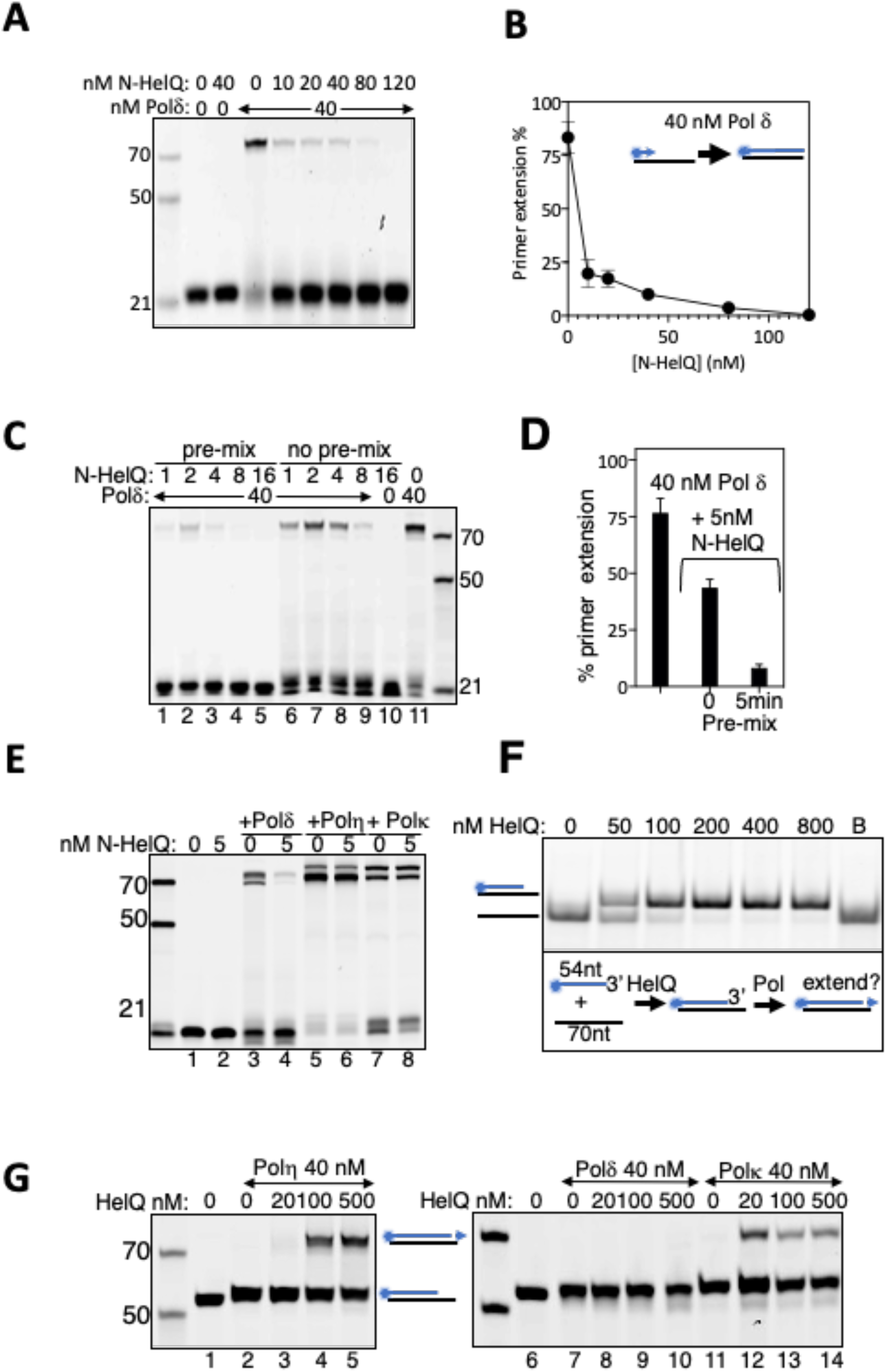
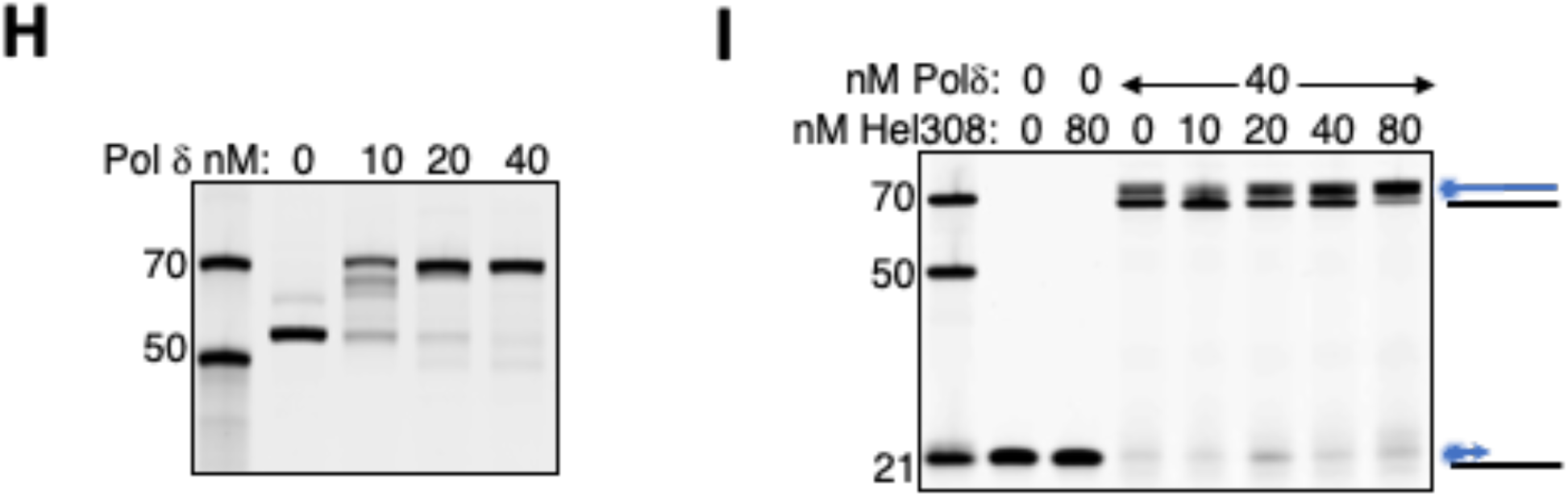
**(A)** DNA synthesis by Pol δ (40 nM, lane 3) was inhibited by titration of N-HelQ as indicated, to zero primer extension when N-HelQ was equimolar (40 nM) with Pol δ, shown in **(B)** where n=3, and showing standard error from mean. **(C)** DNA synthesis by Pol δ (lane 11, 40 nM) was inhibited by N-HelQ (2, 5, 10, 20, 40 nM of N-HelQ) most effectively by pre-incubation with Pol δ for 5 minutes prior to addition of DNA (lanes 1-5), compared with adding the same concentration of proteins separately to DNA simultaneously (lanes 6-10), also in **(D)** quantifying this effect in assays containing Pol δ (40 nM) and N-HelQ (5nM) with either pre-mixing for 5 minutes with prior to adding DNA, or not. N=3, showing standard error from mean. **(E)** DNA synthesis by Pol δ (60 nM) was inhibited by pre-mixing for 5 minutes with N-HelQ (5 nM) (lanes 3 and 4) an effect not observed in the same assays using the other polymerases indicated (lanes 5-14). **(F)** Native acrylamide (TBE) gel showing HelQ (0 - 800nM as indicated) annealing a 70nt DNA strand with a complementary cy5-end labelled 54nt DNA strand (both 15 nM) to generate a recessed 3’ DNA end that can be potentially extended by a polymerase, as shown in the scheme below the gel. Lane B is boiled duplex to release the cy5 end labelled ssDNA. **(G)** Denaturing (urea) gels summarising that the HelQ DNA annealing products from panel F are substrates for active primer extension by purified human DNA polymerases η and κ (lanes 1-5 and 11-14) but not Pol δ (lanes 3-5), despite **(H)** Pol δ extending the same substrate annealed using heat instead of HelQ. **(I)** The archaeal homologue of HelQ lacks the N-HelQ region and is correspondingly unable to inhibit Pol δ in primer extension reactions at concentrations (indicated) effective for HelQ or N-HelQ.

We next checked if selectivity for targeting Pol δ by N-HelQ correlated with similar inhibition from catalytically active full HelQ protein. To do this we generated an assay based on the recently discovered ability of HelQ to anneal DNA strands [25]. In our assays HelQ annealed DNA strands of unequal length (54 and 70 nucleotides), creating a recessed duplex 3’ end with potential for extension by DNA synthesis (Figure 2F). Annealing reactions, and controls lacking HelQ but containing DNA, were inoculated without de-proteinising into primer extension buffer containing Pol η, Pol κ or Pol δ (80 nM). Only Pol δ failed to generate primer extension products from HelQ-annealing reactions (Figure 2G, compare lanes 1-5 and 11-14 with lanes 6-10), despite extending the DNA primer after it was annealed by heating-cooling instead of by HelQ (Figure 2H). We conclude that HelQ selectively inhibits DNA synthesis by Pol δ, requiring its 241 amino acid N-HelQ region – further support for this was observed when the Hel308 helicase-annealase, a close sequence homologue of the HelQ catalytic domains but which lacks the N-HelQ region [31], did not inhibit DNA Pol δ (Figure 2I).

### Mapping interaction of HelQ with Pol δ

Having established that HelQ and N-HelQ both inhibit Pol δ we generated mutant N-HelQ proteins to try to map residues that are required for inhibition. We targeted the two major identifiable parts of N-HelQ – a predicted PWI-like protein fold (amino acids 128 – 237) that utilizes Asp-141 and Phe-142 to disrupt RPA-ssDNA complexes (Figure 3A, RPAi-PWI) [27], and a poorly conserved intrinsically disordered region of N-HelQ (approx. amino acids 1-89). In reactions mixing Pol δ with N-HelQ mutants for 5 minutes prior to adding DNA, mutation of Asp-141 and Phe-142 (N-HelQ^DF-A^) inhibited Pol δ similarly to unmutated N-HelQ (Figure 3B). In agreement with this, truncated N-HelQ lacking the RPA-interacting PWI fold (N-HelQ^ΔRPAi^), therefore comprising only amino acids 1-76 (Figure 3A), was fully proficient at inhibiting Pol δ (Figure 3B). These 76 amino acids are very poorly conserved in HelQ proteins across species except for a tract of basic amino acids that show some conservation (in human HelQ, arginines-51, −52, and −53, and lysine-54, Figure 3A and S2C). Changing all three arginines to glycine (N-HelQ^RGx3^) resulted in a modest but reproducible decrease in inhibition of Pol δ in end point assays (Figure 3B). In the same reactions as a function of time— pre-mixing N-HelQ^RGx3^ with Pol δ from 30-300 seconds prior to adding DNA—gave substantially less inhibition of Pol δ compared with unmutated N-HelQ (Figure 3C and 3D). Therefore, this mutation reduced efficiency of inhibition of Pol δ but without restoring full primer extension activity. It is consistent with these 76 HelQ amino acids forming the site for functional interaction.

**Figure 3:**
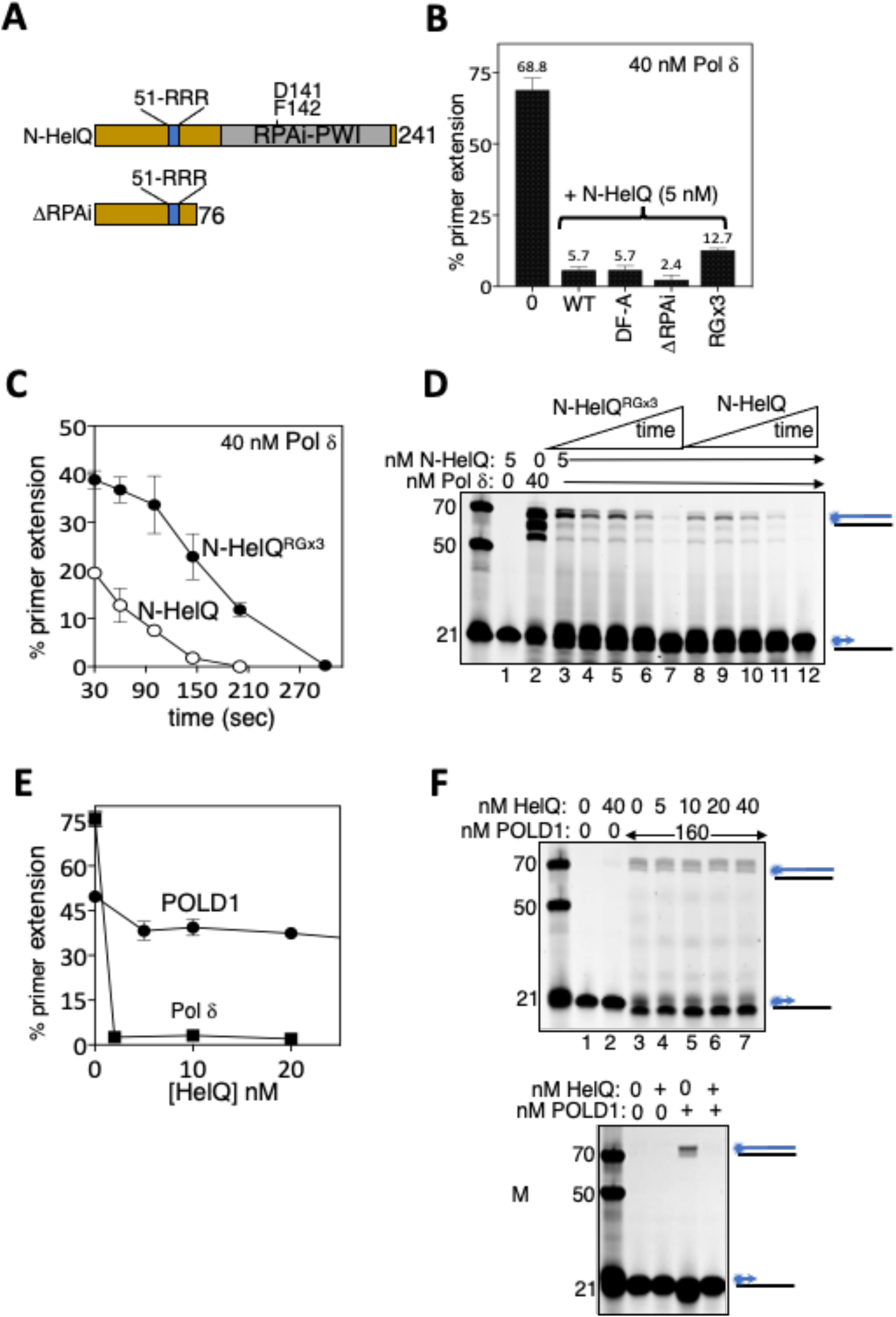
**(A)** Map of the N-HelQ mutant forms tested for inhibition of DNA synthesis by Polδ, see also Figure S2A. Residues R51-R52-R53 in the intrinsically disordered region of N-HelQ were mutated alongside D141 and F142 that are located in a predicted PWI-like fold and are required to remove RPA from ssDNA [27]. The 76-amino acid N-HelQ^ΔRPAi^ fragment was fully proficient at inhibiting Pol δ. **(B)** End point assays measuring inhibition of DNA synthesis by Pol δ (40 nM, maximally in these assays 68.8% of DNA as product) caused by ‘wild type’ N-HelQ (5.7%) compared with mutants as indicated (‘DF-A’, N-HelQ D142AF143A; ΔRPAi, N-HelQ 76 amino acid fragment; RGx3, N-HelQ R51G/R52G/R53G; each at 5 nM). N-HelQ^RGx3^ protein showed comparatively reduced inhibition of Pol δ in these assays (12.7%) (n=3, showing standard error from the mean). **(C)** Time course assays measuring inhibition of DNA synthesis by Pol δ (40 nM) after pre-mixing with N-HelQ or N-HelQ^RGx3^ for 30 seconds to 5 minutes, before adding DNA substrate and dNTPs. The graph shows data points with standard error of mean (n=3). One of the urea gels gels imaged for this data is shown in **(D)**. **(E)** Titration of HelQ (0, 2, 10, 20 nM) did not substantially inhibit DNA synthesis by the isolated POLD1 (160 nM) catalytic subunit of Pol δ, compared with as-expected halting of DNA synthesis by Pol δ (40 nM). **(F)** shows representative imaged urea gels of uninhibited POLD1 product formation after addition of HelQ to 40 nM as indicated. **(G)** Summary urea gel showing that HelQ (40 nM) inhibits primer extension by POLD1-D2-D3 complex (i.e. Pol δ lacking POLD4) (160 nM).

To learn more about how HelQ inhibits primer extension by Pol δ we purified individually the Pol δ catalytic subunit POLD1 (Figure S1A), which synthesises DNA from the short primer without the POLD2-4 subunits of Pol δ (lane 3 in each of Figure 3F). Intriguingly, neither HelQ or N-HelQ inhibited POLD1 (Figure 3E, 3F and S3A) – primer extension products of POLD1 (160nM) were maintained at 49% of the total DNA present, compared to controls of Pol δ complex that were inhibited to zero as expected (Figure 3E). Consistent with this, N-HelQ fully inhibited DNA synthesis by a POLD1-D2-D3 complex (i.e. lacking POLD4) (Figure 3G) – these assays indicate that HelQ does not specifically target the catalytic active site of Pol δ, but instead halts DNA synthesis by targeting POLD2-POLD3. We next tested if any of the individually purified subunits of Pol δ modulate HelQ catalytic activities.

### The POLD3 subunit of Pol δ stimulates DNA annealing and inhibits DNA unwinding by HelQ

Each Pol δ sub-unit (POLD1, D2, D3, and D4) was purified individually (Figure S1A) and tested for modulation of DNA annealing and unwinding by HelQ. HelQ protein (62.5-500 nM) was effective at annealing a pair of complementary 70-nt DNA strands measurably by FRET in real time (0-20 minutes, Figure 4A), and observed in gel-based assays (Figure S3B). Using 125 nM HelQ we repeated the FRET measurements in parallel with reactions additionally containing Pol δ or individual POLD subunits. HelQ alone gave FRET efficiency that increased from 0.11 (time zero) to maximally 0.50 after subtracting values from reactions lacking HelQ (shown in Figure 4B and all panels). Repeat reactions adding Pol δ complex (150nM) alongside the second DNA strand stimulated annealing, increasing FRET efficiency from 0.51 for HelQ alone to 0.76 (Figure 4B). Control reactions containing only Pol δ complex also showed significant annealing maximally to 0.14 (Figure 4B), although this did not account for increased annealing by HelQ. DNA annealing by HelQ was much enhanced by when POLD3 subunit alone was added to HelQ reactions – POLD3 increased FRET efficiency from 0.50 to 0.87, with only very weak annealing (maximally 0.07) observed for POLD3 alone (Figure 4C). Stimulation by POLD3 contrasted with no significant effect observed in reactions adding subunits POLD1, POLD2 or POLD4 (Figures 4D-F).

**Figure 4.**
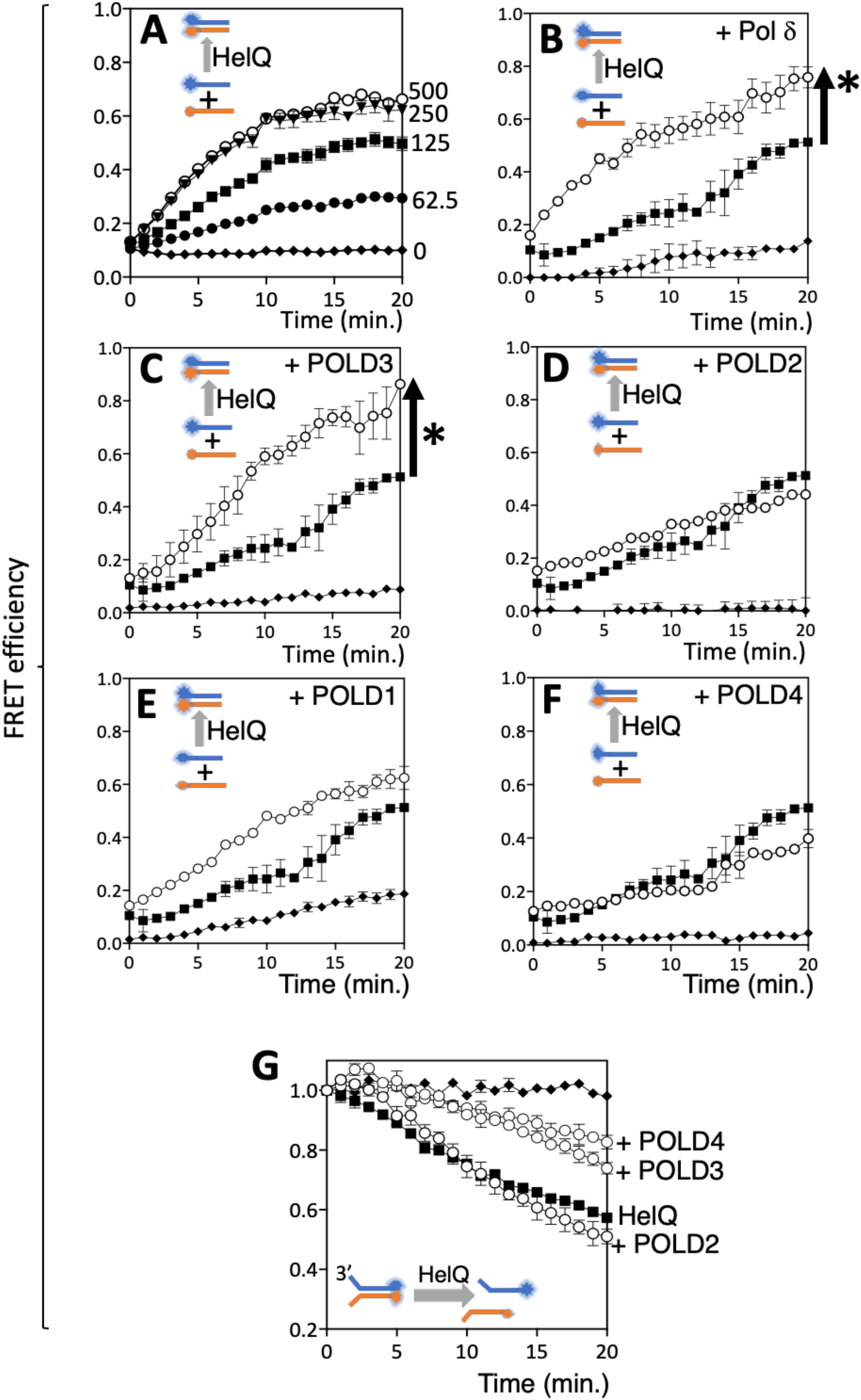
**(A)** Real-time FRET measurements of annealing two Cy5/Cy3 end-labelled complementary DNA strands by HelQ at the nM concentrations indicted. **(B) – (F)** FRET measurements of DNA annealing by HelQ alone (125 nM, black squares) compared with addition of Pol δ, POLD1, POLD2, POLD3 or POLD4 (150 nM, in each panel white circles) to HelQ annealing reactions as indicated. Each plot also shows the FRET measurement for reactions lacking HelQ (black diamonds) therefore containing only Pol δ or POLD1-4, after adjusting for zero protein control reactions. The plots are means of two experiments showing error bars representing standard deviation from the mean. In each panel HelQ annealing is The arrow and * next to panels B and C highlight the only significant (compared with controls) increase in annealing by HelQ. **(G)** FRET measurements of DNA unwinding by HelQ (125 nM, plotted as black squares) inhibited by addition of POLD3 or POLD4, but not POLD2 (all 150 nM, plotted as white circles). Reactions containing zero protein are denoted by black diamonds. See also Figure S3A that additionally shows the plots for POLD-only reactions, omitted from this figure for clarity. The plots are means of two experiments with error bars showing the standard deviation from the mean

In FRET assays of HelQ helicase activity (Figure 4G) – reactions contained 1 mM ATP and 1 mM MgCl_2_ and a cy3/cy5 DNA fork for strand separation – POLD3 and POLD4 both inhibited DNA unwinding, increasing the FRET efficiency from 0.57 for uninhibited HelQ helicase, to 0.74 and 0.81, respectively (Figure 4G), after adjusting for the control reactions containing no protein (shown in 4G) and only POLD3 or POLD4 (Figure S3C). POLD2 had no effect on HelQ helicase activity (Figure 4G). HelQ helicase assays require Mg^2+^ therefore Pol δ and POLD1 could not be tested for any modulating effect Mg^2+^ activates their exonuclease activity to degrading the helicase substrate with 3’ to 5’ directionality, the same as DNA translocation by HelQ. However, the data show that Pol δ and POLD3 modulate HelQ by stimulating its annealing activity and inhibiting DNA unwinding.

## Discussion

HelQ is a metazoan-specific protein implicated in maintaining genetic stability across multiple DNA break repair pathways [21, 22, 24, 25]. Here we demonstrate that HelQ is a potent inhibitor of DNA synthesis by Pol δ, and that Pol δ and its isolated POLD3 subunit stimulate DNA single strand annealing by HelQ. These functional interactions provide a plausible explanation for loss of HelQ from cells (*HELQ*^*-/-*^) increasing tandem DNA duplications and long tract recombination. Previous work [25, 27] has shown that HelQ removes RPA from single-stranded DNA, and that ssDNA triggers its DNA translocase activity [27, 29], that is further stimulated by Rad51 [25]. Alongside the data presented in this work a model emerges for HelQ as a controller of homologous recombination by assembling onto ssDNA *via* RPA, for ATP-dependent DNA translocation until it encounters Pol δ, ending DNA synthesis and annealing DNA strands, summarised in Figure 5.

**Figure 5.**
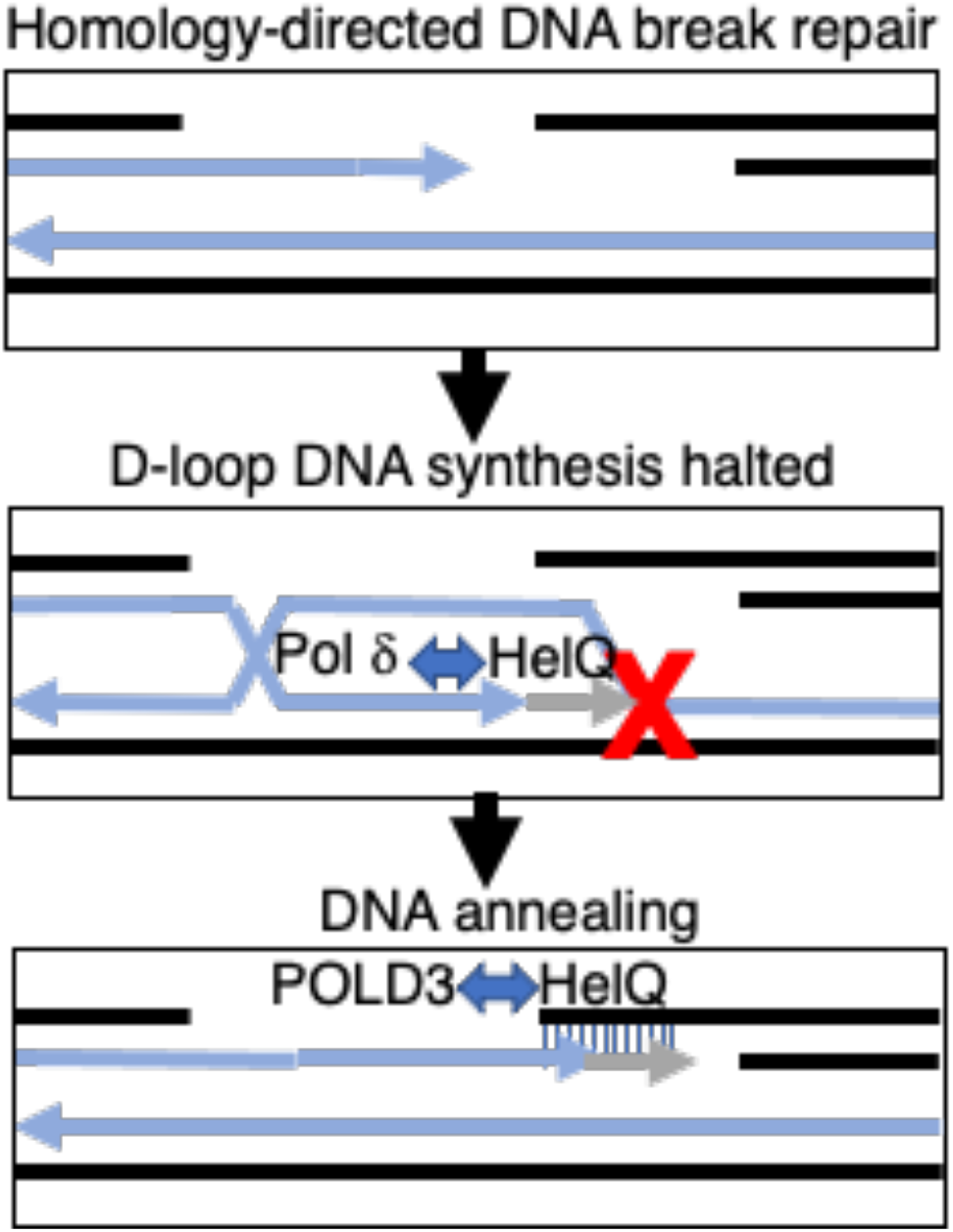
HelQ as a controller of multiple modes of homology-directed DNA repair in metazoans, by inhibiting Pol δ and triggering DNA annealing. This is proposed to limit the extent of replicative homologous recombination by ending DNA synthesis and promoting DNA single-strand annealing.

The 76 amino acid fragment at the extreme N-terminal end of HelQ (N-HelQ 1-76) was fully proficient at inhibiting Pol δ, an effect diminished by mutating a tract of arginine residues in this fragment. In contrast, mutation of N-HelQ residues that are critical for RPA/N-HelQ interaction (Asp-141 and Phe-142) had no effect on Pol δ/N-HelQ functional interaction. Therefore N-HelQ seems to provide a ‘hub’ controlling activities of other proteins by distinct protein-protein interactions. The majority of the 250 amino acid N-HelQ sequence is intrinsically disordered and with little detectable sequence conservation across HelQ proteins in metazoans. The intrinsic disorder of N-HelQ may explain why we have not been able to observe stable physical interaction between Pol δ or POLD3 with HelQ or N-HelQ in any permutations in EMSAs and affinity pull-downs. Several lines of evidence were consistent with functional interaction between POLD3 and HelQ, including stimulation of DNA single strand annealing and the inability of HelQ to inhibit basal level DNA synthesis by POLD1 alone. POLD3 is required, alongside the Pif1 helicase, as a regulatory factor promoting DNA synthesis during human homologous recombination by Break-Induced Replication [32], where it is thought to stabilise the Pol δ complex. POLD3 is also required for DNA synthesis by the REV3L subunit of human polymerase ζ complex that drives mutagenic DNA synthesis in several contexts, including Microhomology-Mediated Break Induced Replication [33]. The recent insights from genetics alongside new biochemical data presented here advances a model for HelQ protein as a restraint to DNA synthesis arising from unproductive homologous recombination and other DNA homology-dependent repair processes.

## Methods

### Protein purifications

Full-length human HelQ and HelQ^ΔWHD^ proteins were generated and purified as described in [27] and see Figure S1A. N-HelQ protein and mutants N-HelQ^RGx3^, N-HelQ^ΔRPAi^ and N-HelQ^DF-A^ were purified as in [27], except for additional inclusion of a heparin column step, as described for HelQ. N-HelQ proteins did not bind to heparin, collecting in flow-through fractions. Purification of human DNA polymerase δ complex (PolD1-4) was as described in [28], and DNA polymerase κ (PolK) as in [34].

Human full-length DNA polymerase η (PolH) was expressed in *E. coli* strain BL21 (DE3) with an N-terminal (His)_6_-SUMO tag from a pE-SUMO-pro expression vector (LifeSensors). Cells were grown in 2YT media at 24°C to an OD_600_ of 0.8 and then induced with 0.1 mM isopropyl β-D-thiogalactopyranoside (IPTG) and incubated further for 19 hrs at 16o C. Cells were collected by centrifugation and re-suspended in lysis buffer [50 mM Tris pH (8), 750 mM NaCl, 40 mM imidazole, 5 mM β-Mercaptoethanol, 0.2% NP-40, 1mM PMSF, 5% Glycerol and EDTA free protease inhibitor cocktail tablet/50ml (Roche, UK)]. All further steps were performed at 4o C. Cells were lysed enzymatically by adding 2 mg/ml lysozyme and mechanically by sonication. Cell debris was removed by centrifugation (22,040 g, 60 min) and the clear supernatant was directly loaded onto HisTrap HP 5 ml affinity column (Cytiva) pre-equilibrated with buffer A (50 mM Tris (pH 7.5), 500 mM NaCl, 40 mM imidazole, 5 mM β-Mercaptoethanol and 5% Glycerol). The column was then washed with 50 ml of buffer A followed by elution gradient with 50 ml of buffer B (50 mM Tris (pH 7.5), 500 mM NaCl, 500 mM imidazole, 5mM β-Mercaptoethanol and 5% Glycerol). The protein was eluted at ∼310 mM imidazole. The peak fractions were pooled and dialyzed overnight in dialysis buffer (50mM Tris (pH 7.5), 500 mM NaCl, 5 mM β-Mercaptoethanol and 5% Glycerol) in the presence of SUMO protease (LifeSensors) to remove the SUMO tag to generate native PolH. The dialyzed sample was loaded again onto HisTrap HP 5ml using buffers A and B and un-tagged PolH was collected in the flow-through fractions. Fractions that contained PolH were collected, concentrated, and loaded onto HiLoad 16/600 Superdex 200 pg (Cytiva) equilibrated with storage buffer (50 mM Tris (pH 7.5), 300 mM NaCl, 10% Glycerol and 1 mM DTT). Fractions containing PolH were concentrated, flash-frozen and stored at −80 °C.

The human Pol δ sub-units POLD1, POLD2, POLD3, and POLD4 were expressed individually in *E. coli* BL21 A.I., each with an N-terminal (His)_6_ tag from plasmids generated by GeneArt (ThermoFisher Scientific). For each purification, cells were grown in LB broth with 100 μg/mL ampicillin at 37 °C until OD_600_ reached 0.6 when IPTG, and L-arabinose were added to 0.5 mM and 0.2 % (w/v), respectively. Growth was continued for 18 h at 20°C before harvesting cells for storage at −80 °C in 50 mM Tris-HCl pH 8.0, 1 M NaCl, 5 % (v/v) glycerol and 0.5 mM phenylmethylsulphonyl fluoride (PMSF). Each sub-unit was purified individually after thawing cells for sonication, and clarification as soluble proteins POLD1, POLD2, POLD3, or POLD4. Each was passed into a Ni-NTA column (GE Healthcare) pre-equilibrated with buffer A (50 mM Tris-HCl pH 8.0, 10 mM imidazole, 1 M NaCl, 5 % (v/v) glycerol). Bound protein was eluted using this buffer containing 500 mM imidazole. POLD1 was concentrated to 1 mL using Vivaspin centrifugal concentrator (Sartorius) and loaded onto a HiLoad 16/600 Superdex 200 pg column (GE Healthcare) pre-equilibrated with 50 mM Tris-HCl pH 8.0, 100 mM NaCl, 5 % (v/v) glycerol. Eluted POLD1 was passed into a 1 mL heparin column (GE Healthcare) pre-equilibrated using the same buffer—POLD1 eluted from heparin column in the same buffer at 1 M NaCl, and pooled fractions were dialyzed into buffer Tris-HCl pH 8.0, 200 mM NaCl and 20 % (v/v) glycerol for storage at −80 °C. POLD2 after Ni-NTA and dialysis was further purified through binding to a heparin column, using the same procedure as for POLD1, and eluted POLD2 was finally passed through a Superdex 16/600 column before dialysis and storage as for POLD1. POLD3 was purified as for POLD1 and POLD2 initially (sequential Ni-NTA and heparin columns) except that it did not bind to heparin and was loaded directly into a 5 mL HiTrap Q-Sepharose column to which it bound and was eluted from 300-600 mM NaCl in otherwise the same buffer as for heparin. POLD3 fractions were then passed through a Superdex 16/600 column before dialysis and storage as for POLD1 and POLD2. POLD4 was pure to apparent homogeneity after a Ni-NTA column and was dialysed and stored as for the other sub-units. Figure S1A shows each final purified protein. *E. coli* DNA polymerase I was purchased from New England Biolabs. *E. coli* DNA polymerase III was a kind gift from Dr. Michelle Hawkins, University of York, UK.

### DNA substrates

M13 single-stranded DNA was purchased from New England Biolabs. DNA substrates are detailed in the Supplementary data (Table S1). Oligonucleotides (Sigma-Aldrich) were 5’ Cy5 end labelled for DNA annealing and primer extension reactions, and additionally with Cy3 for FRET-based DNA helicase assays. DNA substrates were annealed by heating a 1.2:1 ratio of unlabelled to Cy5-labelled oligonucleotides to 95oC for 10 minutes and annealing to room temperature overnight. Annealed DNA was separated from unannealed oligonucleotide by gel electrophoresis through 10% (w/v) acrylamide Tris.HCl-borate-EDTA (TBE) gels, followed by excision of the desired substrate as a gel slice gel and soaking overnight into Tris.HCl pH 7.5 buffer containing 150 mM NaCl to recover DNA from gel pieces.

### Primer extension assays

Primer extension reactions (25 μl) were based on [35, 36], utilizing purified human Pol δ (40 nM) in a holoenzyme complex with PCNA (40 nM), RFC (10 nM) and RPA (320 nM) for 30 minutes at 37oC before stopping the reactions by adding 2 μl of stop solution (50 mM Tris pH 8.0, 100 mM EDTA, 0.1% w/v SDS and 5 mg/ml of proteinase K) and observing the products after electrophoresis through a 0.8% agarose TBE gel. RPA, PCNA and RFC were pre-mixed for 10 minutes on ice in the reaction buffer containing DNA substrate (82.5 ng M13-primer DNA/reaction) and dNTPs, before adding Pol δ to start the reactions.

Primer extension reactions by isolated Pol δ were in 20 μl containing substrate DNA (21 nt primer oligo annealed to a 70 nt oligo template, each at 15 nM), 10 mM MgCl_2_, 40 mM Tris-HCl pH7.5, 1 mM DTT, 0.2 mg/mL BSA, 50 mM NaCl and 200 μM of each dNTP. Primer extension assays using human DNA polymerase η and polymerase k were in buffer 40 mM Tris-HCl pH 8.0, 10 mM DTT, 0.25 mg/mL BSA, 60 mM KCl, 2.5 % (v/v) glycerol, 5 mM MgCl_2_, 200 μM dNTPs. Primer extension by *E. coli* Pol III core (DnaE) and Polymerase I were in buffer 10 mM magnesium acetate, 40 mM HEPES-NaOH pH 8.0, 0.1 mg/mL BSA, 10 mM DTT and 200 μM dNTPs. Unless stated otherwise, all polymerase primer extension assays were for 30 minutes at 37°C after adding DNA. Reactions were halted by adding 5ml of stock stop solution, comprising 50 mM Tris pH 8.0, 100 mM EDTA, 5 mg/mL proteinase K and 1% (w/v) SDS, and allowing room temperature incubation for 2-5 minutes. Stopped reactions were mixed with loading dye containing 20% (v/v) glycerol, 78% (v/v) formamide and Orange G for electrophoresis through 10% (w/v) acrylamide TBE denaturing (8 M urea) gels, at 5 watts for 3 hours. Primer extension products were visualised *via* the Cy5-DNA end label using a Typhoon scanner, and files were quantified using ImageJ and Prism software. For primer extension reactions containing HelQ, proteins were pre-mixed in their storage buffers, in isolation from reaction buffer and DNA, and reactions commenced by adding buffer containing DNA and dNTPs.

### DNA annealing assays

DNA annealing assays contained DNA strands detailed in Supplementary S1. For gel-based assays, reaction mixtures contained 20 mM Tris.HCl pH 7.5, 100 mg/mL BSA, 7% (v/v) glycerol, 25 mM DTT, DNA strand 1 (15 nM). HelQ was added for incubation for 2 minutes at room temperature, and then annealing began by addition of 15nM of DNA strand 2, for 5 minutes at 37°C when reactions were halted by adding 5ml of stop solution. Annealing products were assessed in 10% acrylamide (w/v) TBE gels electrophoresed for 1 hour at 150 volts.

For FRET assays, reaction mixtures contained 20 mM Tris.HCl pH 7.5, 100 mg/mL BSA, 7% (v/v) glycerol, 5 mM DTT, and 5’Cy5 labelled ELB-41 (50 nM), and with addition of 1 mM ATP and 1 mM magnesium chloride if stated. HelQ was added for incubation for 2 minutes at room temperature, and then annealing began by addition of Cy3 labelled ELB40 (50 nM). Fluorescence was measured on a 37oC pre-incubated FLUOstar Omega (BMG Labtech) at excitation of 540nm and emissions at 590nm and 670nM accounting for background Cy3 fluorescence and FRET pair excitation respectively. Readings were taken every minute for 20 minutes. Annealed DNA was calculated by normalising the corrected data to a fully pre-annealed control (representing 100% annealing).

### DNA helicase assays and EMSAs

Helicase unwinding assays measured separation of annealed DNA strands by monitoring the migration of Cy5 labelled single-stranded DNA. Reaction mixtures contained 20 mM Tris.HCl pH 7.5, 100 mg/mL BSA, 7% (v/v) glycerol, 25 mM DTT, 5 mM ATP, 5mM magnesium chloride and 25 nM Cy5 end-labelled DNA fork substrate and protein concentrations indicated. Reactions were incubated at 37oC for 10 minutes prior to quenching by addition of stop buffer. Reactions were resolved on 10% (w/v) acrylamide TBE gels and analysed using an Amersham Typhoon phosphor-imager. Percentage unwinding was calculated by determining the percentage of Cy5 labelled ssDNA compared with Cy5 labelled forked DNA.

For real-time FRET assays, reaction mixtures contained 20 mM Tris.HCl pH 7.5, 100mg/mL BSA, 7% (v/v) glycerol, 5 mM DTT, 5 mM ATP, 5 mM magnesium chloride and 50 nM FRET pair labelled DNA fork substrate. On addition of protein, fluorescence was measured in a 37oC pre-incubated FLUOstar Omega (BMG Labtech) at excitation of 540 nm and emissions at 590 nm and 670 nm accounting for, respectively, background Cy3 fluorescence and FRET pair excitation. Readings were taken every minute for 20 minutes, with undissociated substrate calculated by normalising the corrected data to time 0 min.

Electrophoretic mobility shift assay (EMSAs) reaction mixtures contained 20 mM Tris.HCl pH 7.5, 100mg/mL BSA, 7% (v/v) glycerol, 25 mM DTT, 50 mM EDTA and 25 nM fluorescently labelled DNA and varying concentrations of HelQ and variant mutant proteins. Assembled complexes were incubated at 37oC for 10 minutes prior to resolution on 5% (w/v) acrylamide TBE gels in Orange G loading dye. Gels were imaged and analysed using an Amersham Typhoon phosphor-imager to detect migration of Cy5 end-labelled DNA.

## Acknowledgments

The work was supported by Nanna Therapeutics (Cambridge, U.K.) to ELB, The BBSRC (grant number BB/T006625/1 to ELB and grant number BB/R013357/1 to PS), King Abdullah University of Science and Technology Baseline Funding BAS/1/1002-01-01 to SMH, and PhD studentships from The BBSRC DTP and the University of Nottingham School of Chemistry. We thank Stuart Wood, Aurelio Reyes, Christopher Cooper and Stan Jozwiakowski for helpful comments on the work.

## Author Contributions

ELB, LH and RL designed experiments, and generated and interpreted the data. LH, RL, MT, AH, AON, AC, TJ and MF acquired data and generated constructs and/or proteins. ELB wrote the manuscript, and PS and SH interpreted data and contributed to writing the manuscript.

---

**Supplementary Table S1.**
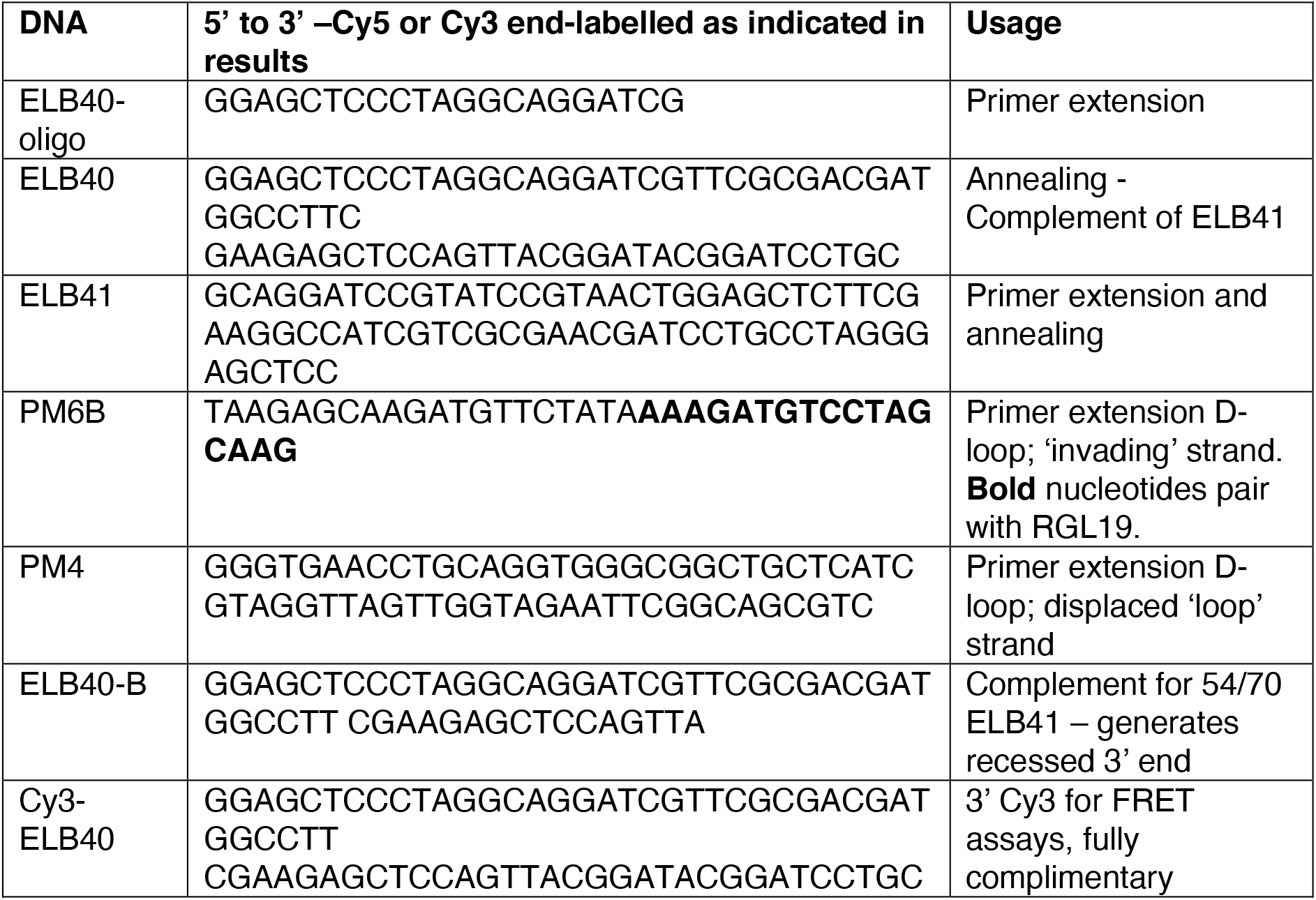
DNA substrates used in this work for DNA binding EMSAs, anisotropy, primer extension and DNA annealing reactions

## Supplementary Figures

**Figure S1A.**
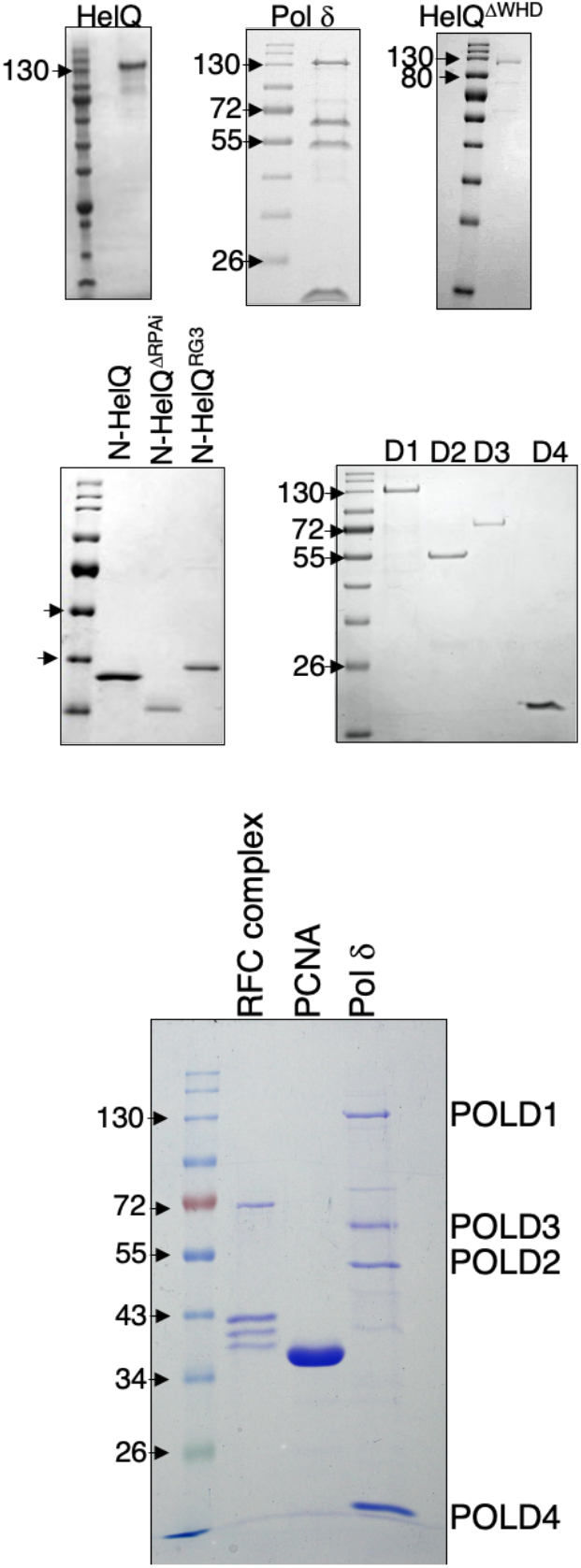
Coomassie stained SDS-PAGE gels of purified proteins used in this work, alongside RPA protein shown in [1]. In all panels protein size markers are shown.

**Figure S1B.**
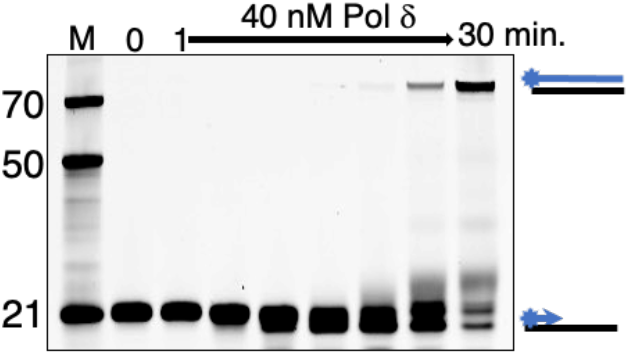
Urea gel summarising time course (0, 1, 2, 5, 10, 15, 20 and 30 minutes) DNA synthesis by Pol δ (40 nM) observed as extension of a 21 nucleotide (nt) cy5-labelled DNA annealed to a 70 nt template. Known length cy5 DNA markers are shown to the left of the panel, and in subsequent panels.

**Figure S1C.**
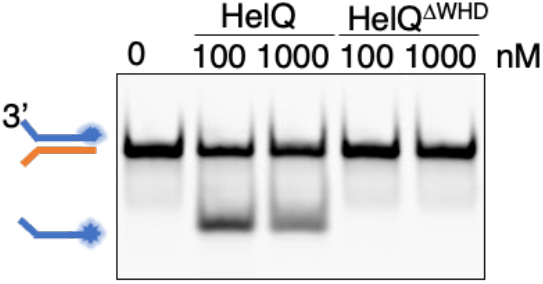
Native TBE gel showing helicase unwinding products from a DNA fork substrate (15 nM) of HelQ compared with HelQ^ΔWHD^ at concentrations shown.

**Figure S1D.**
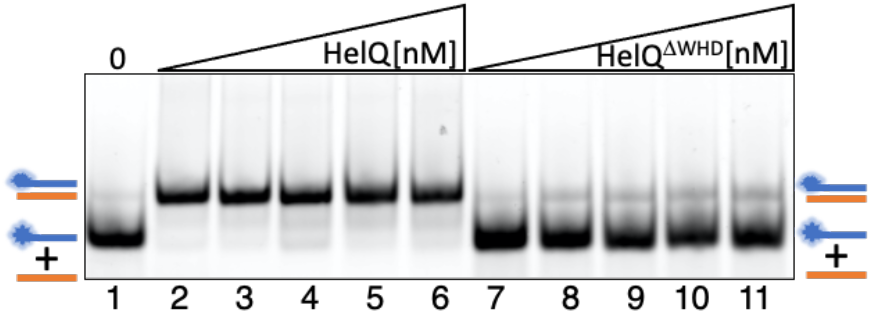
Native TBE gel showing HelQ or HelQ^ΔWHD^ DNA single strand annealing products from two complementary 70 nucleotide strands (each 15 nM). Protein concentrations for this assay were 200, 400, 600, 800 and 1000 nM to test the mutant HelQ to the maximum concentration possible.

**Figure S2A.**
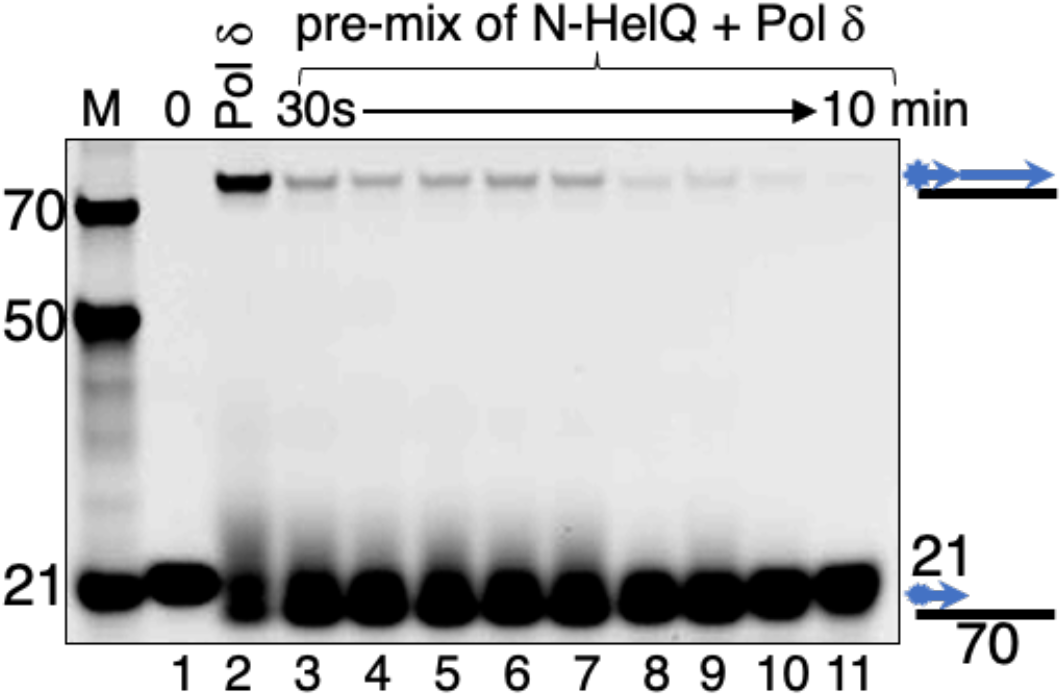
Urea gel showing primer extension by Pol δ (40 nM) in 30-minute reactions when in the absence of pre-mixing with N-HelQ (lane 2) or after pre-mixing with N-HelQ (10 nM) for periods of time as indicated; 30, 60, 90, 120, 180, 240, 300, 420 and 600 seconds (lanes 3-11) prior to adding DNA to the proteins.

**Figure S2B.**
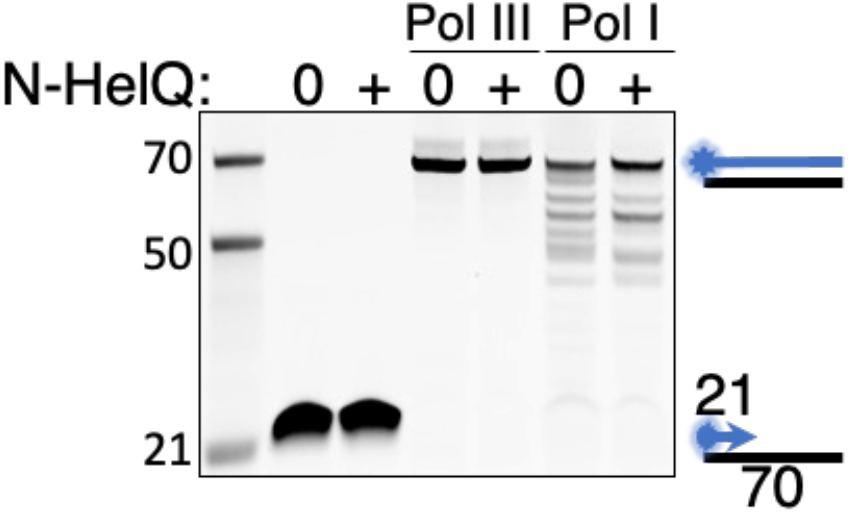
Urea gel summarising that N-HelQ (10 nM) has no effect on DNA synthesis by primer extension catalysed by *E. coli* DNA polymerases PolI and PolIII (each 80 nM), with DNA length markers as indicated.

**Figure S3A.**
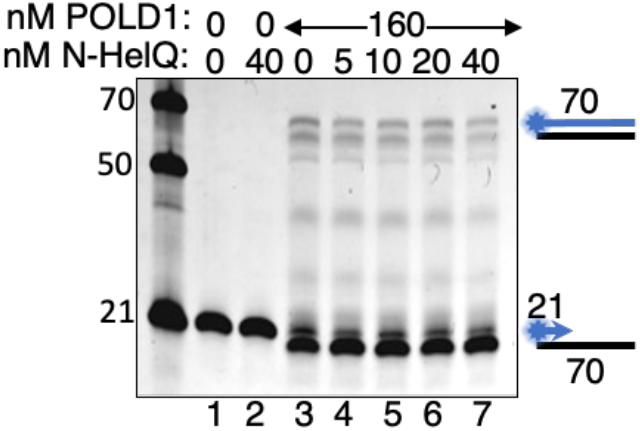
Urea gel summarising that basal-level primer extension by POLD1 (lane 3, 160 nM as indicated) is unaffected by titration of N-HelQ as indicated, in agreement with data in Figure 3 using full HelQ.

**Figure S3B.**
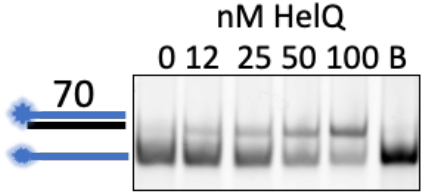
Native TBE gel summarising DNA annealing by HelQ (concentrations as indicated) corresponding to FRET measurements.

**Figure S3C.**
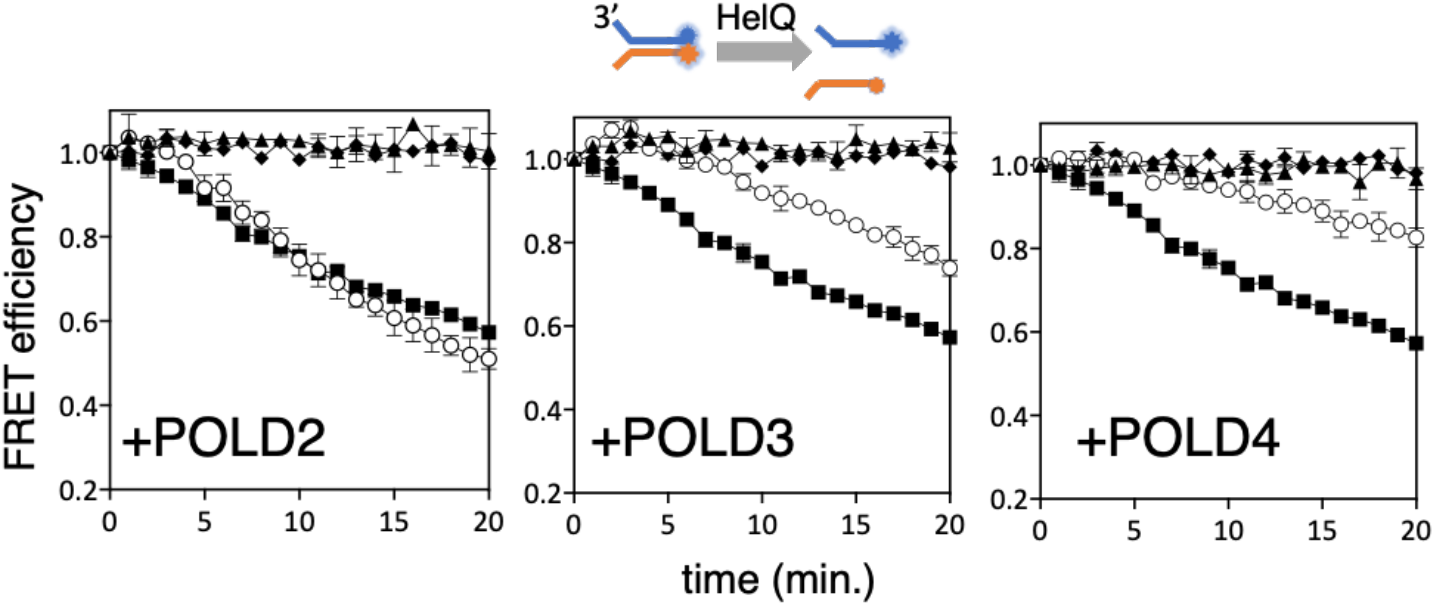
Plots of FRET measurements for DNA unwinding by HelQ only (black squares), and after addition of POLD2, POLD3 or POLD4 (white circles, as indicated in each panel), compared with plots for POLD2, POLD3 or POLD4 only (black triangles, as indicated in each panel). The plots of black diamonds are for zero protein control reactions.

